# Biological oscillations without genetic oscillator or external forcing

**DOI:** 10.1101/2024.09.20.614027

**Authors:** Vincent Vandenbroucke, Lucas Henrion, Frank Delvigne

## Abstract

Oscillators are fundamental to biological systems, underpinning essential processes such as cell division, circadian rhythms, and developmental cycles. While both natural and synthetic genetic oscillators have been extensively studied, oscillatory behaviors in cells can also emerge without dedicated genetic circuits. In earlier work, we uncovered sustained oscillations in phenotypic switching across diverse cellular systems and gene circuits, occurring spontaneously, without external forcing and linked them to the induction of slow-growing phenotypes. In this study, we identify the conditions that give rise to such intrinsic phenotypic instabilities, leading to population-level oscillations. We develop and analytically solve a simplified mathematical model of a stress-induced phenotype, mapping the range of continuous culture conditions that trigger oscillatory gene expression. This instability range, predicted by the model, was experimentally validated in *Bacillus subtilis* cultures. Our findings reveal that oscillations can arise in the complete absence of genetic oscillators or external perturbations. Although demonstrated here for a stress response in continuous culture, this phenomenon may occur in any long-term cultivation where environmental feedback links an inducer to the cellular system, broadening the landscape of possible oscillatory behaviors in microbiology and synthetic biology.

## Introduction

Oscillations in gene expression are traditionally attributed to dedicated genetic circuits, metabolic pathways or external forcing^1 5^. Here, we report sustained oscillations emerging in the complete absence of either, arising instead from the interplay between cellular burden and nutrient availability in a constant environment. Oscillatory behaviors are pervasive in biology, spanning multiple scales, from single cells to entire organisms, and regulating substantial portions of cellular metabolism^2 6^. Their study has revealed natural gene circuits acting as biological oscillators, such as the circadian clock in cyanobacteria^8^ and gut microbiota^9 12^, as well as the cell cycle in *E. coli* ^14^. These discoveries have been complemented by synthetic designs^16 18^, including the iconic repressillator, which generates spontaneous oscillations in gene expression through a triad of repressor proteins^19 23^. In general, biological oscillations arise from dedicated gene circuits ^23 16^, external periodic forcing^24^, or a combination of both^25^. In all reported cases, oscillations under constant culture conditions were linked to an underlying genetic oscillator e.g., yeast respiratory cycles^3^, bacterial cell cycle^7^ and plankton synchronization^4^.

Surprisingly, we previously observed that highly burdensome gene circuits can exhibit sustained oscillations in a simple chemostat^10^, a continuous cultivation device initially designed to maintain constant growth conditions^26 27^. Strikingly, the two systems that displayed this behavior, the T7 expression system in *E. coli* BL21 and the sporulation program of *Bacillus subtilis*, lack network architectures typically associated with oscillatory dynamics. To uncover the origin of these instabilities, we focused on the *B. subtilis* system, which can be distilled into: i) Two discrete phenotypes linked to growth capacity, vegetative, non-fluorescent cells that grow, and sporulating, fluorescent cells that do not; ii) A single trigger, glucose, whose depletion both induces sporulation^11^ and serves as the primary carbon source.

To uncover the origin of these unexpected oscillations, we developed a minimal mathematical model of the stress response in a chemostat, a system with constant nutrient influx, where stress is triggered when nutrient levels fall below a critical threshold. The model predicts that, even under the classically “stable” operating conditions of a chemostat, there exists a specific range of dilution rates in which no steady-state population-level stress response can be sustained. This challenges the long-standing assumption that chemostats inherently stabilize gene expression across microbial populations^26^. We experimentally tested this prediction using *Bacillus subtilis* 168 carrying a chromosomal *gfp* reporter under the control of the *spoIIE* promoter, SpoIIE being a key regulator of asymmetric cell division during sporulation^13 15^. Cultures were grown at different dilution rates, and sporulation activation was tracked at the single-cell level using a custom flow cytometry– bioreactor interface^17 20^. Consistent with model predictions, we observed oscillatory sporulation dynamics within the predicted operating range, manifesting as successive bursts of sporulating cells sweeping through the population.

## Results

### A mathematical model predicts instabilities in gene expression for a specific range of dilution rate

To probe how oscillations can arise in the absence of a genetic oscillator, we constructed a minimal ODE model capturing stress-induced dynamics in a cell population. The framework tracks two phenotypic states: non-stressed cells (*X*_1_) and stressed cells (*X*_2_) (**Eqs. 1–5**). The stressed phenotype grows more slowly, with a maximum growth rate *μ*_*max*,2_ = *k μ*_*max*,1_ where *k* ∈ [0, 1] and *μ*_*max*_ is the maximal growth rate of the non-stressed biomass. Switching from *X*_1_ to *X*_2_ occurs at a constant rate *H* when the substrate concentration *S* drops below a threshold *K*_*I*_ and is implemented via a step function *H*_1_*(S)* (**Eq. 5**). Because stressed cells grow slowly and stress-related proteins are primarily diluted through cell growth, the reverse transition from *X*_2_ to *X*_1_ is assumed negligible.

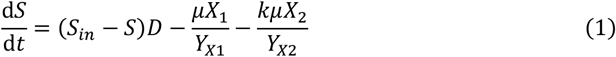

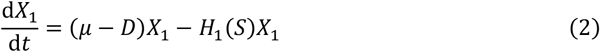

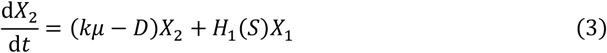

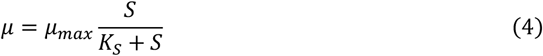

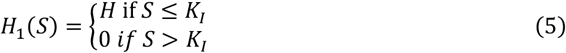

Here, *S*_*in*_ denotes the sugar concentration in the feed, *D* the dilution rate, *Y*_*Xi*_ the biomass yields of each phenotype, and *K*_*S*_ the sugar concentration at half-maximum growth rate. Analytical solutions of the model (see **SI Appendix**) reveal a critical range of dilution rates in which no steady-state equilibrium can be achieved among the state variables, rendering the system inherently unstable (**Fig. 1A**).

**Figure 1:**
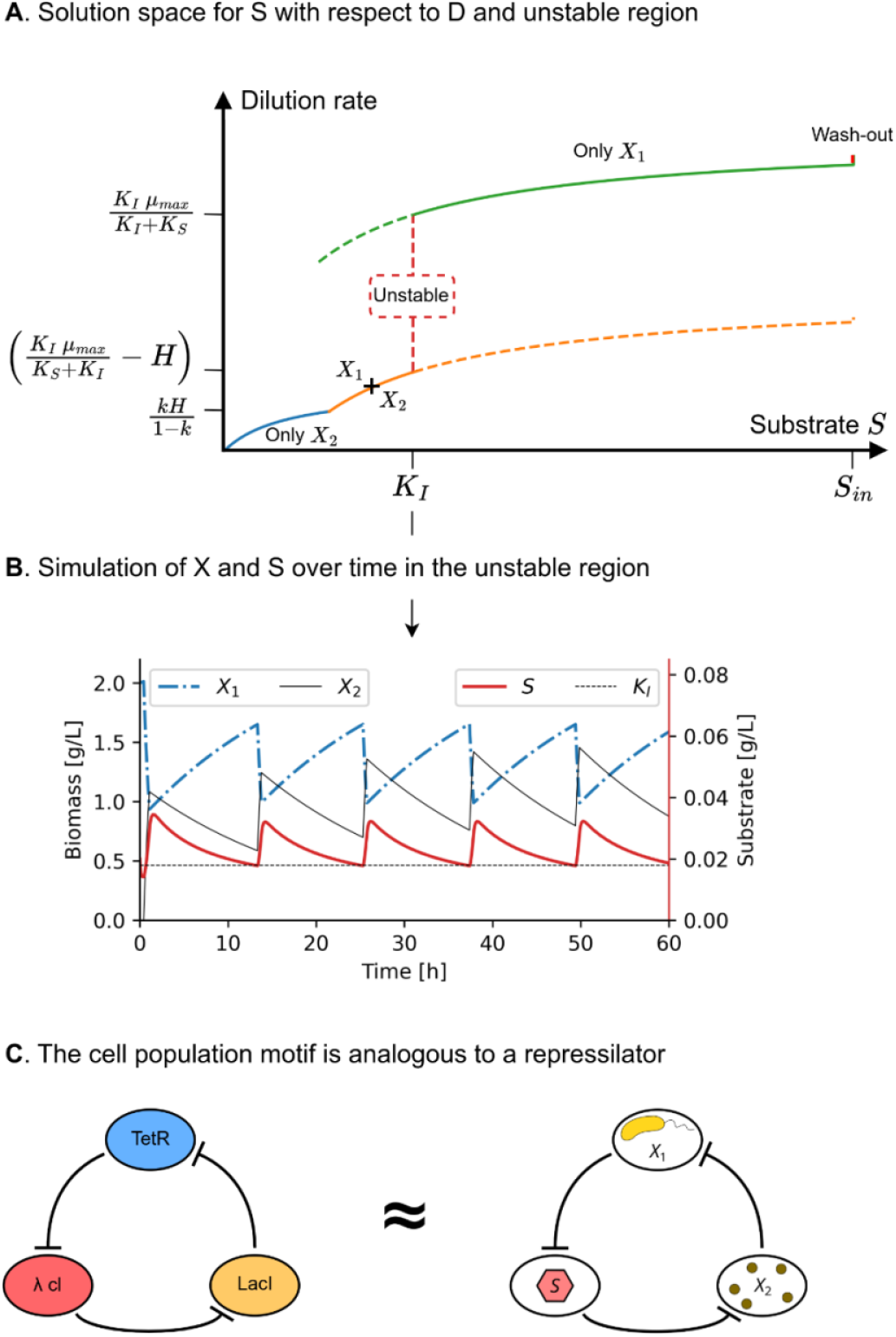
Representation of the solution space of the stress-response model at steady state. **(A)** Model predictions reveal three distinct regimes. At high dilution rates, only the non-stressed phenotype is sustained, as the sugar concentration remains above the threshold *K*_*I*_. At low dilution rates, the stressed phenotype dominates, with sugar concentrations falling below *K*_*I*_. Between these extremes lies an instability window where no steady state exists, and oscillations can emerge. **(B)** Introducing a delay in the stress response generates sustained oscillations *in silico* within the unstable region. The example shown corresponds to *B. subtilis* sporulation dynamics with *D* = 0.1 h^−1^ and a 0.5 h delay, although small parameter changes can markedly alter the oscillatory behavior. **(C)** Schematic analogy between the sporulation system in *B. subtilis* (right) and the synthetic repressilator (7) (left). Here, the stressed and non-stressed subpopulations, together with the extracellular sugar concentration, form a triple negative feedback loop. In the chemostat, oscillations arise at the level of reactor concentrations rather than intracellular protein levels: sporulation (or stress induction) reduces the number of unstressed cells, decreasing sugar uptake, raising sugar concentration above *K*_*I*_, inhibiting sporulation, and allowing unstressed cells to expand again.

Within this instability window, two mutually exclusive conditions prevent convergence to equilibrium. First, the classical chemostat formulation, omitting the stress phenotype, fails because the residual substrate concentration *S* stabilizes below the threshold *K*_*I*_, triggering the stress response. Second, the alternative scenario of continuous switching into the stressed state is also invalid, as the associated reduction in substrate uptake drives *S* above *K*_*I*_, thereby suppressing the stress response.

Rather than resolving into either case, the system hovers near the switching threshold *K*_*I*_, forcing cells to continually adjust to environmental fluctuations. This creates a negative feedback loop on population growth which, in combination with delayed phenotypic transitions, produces sustained oscillations analogous to those of the synthetic repressilator^23 19 28^ (**Fig. 1B,C**). The appearance and persistence of these oscillations are expected to depend on both the delay magnitude and the fidelity of the step function in capturing the switching dynamics, while the dilution-rate range for their occurrence can be experimentally validated.

### The instability region can be experimentally observed in chemostat for a given range of dilution rates

Model predictions were tested under continuous cultivation, a regime designed to maintain stable growth conditions through substrate (*S*) limitation, which can be tuned via the dilution rate (*D*). As a case study, we examined *Bacillus subtilis* sporulation, a stress response triggered when residual glucose concentration falls below a threshold *K*_*I*_. Here, *X*_1_ denotes vegetative (non-stressed) biomass and *X*_2_ denotes stressed biomass (spores). Sporulation was monitored at the single-cell level using automated flow cytometry and a P_*spoIIE*_*::gfp* transcriptional reporter, where *spoIIE* encodes a key regulator of asymmetric division during sporulation.

Using *H, K*_*I*_, *μ*_*max*_, and *K*_*S*_ values determined from batch cultures (**SI Appendix**), and assuming no growth for the stressed phenotype, the model predicts that at high *D*, with residual glucose above *K*_*I*_, only *X*_1_ persists. At low *D*, with residual glucose at or below *K*_*I*_, both *X*_1_ and *X*_2_ are present, with *X*_2_ (spores) periodically increasing in abundance. Due to the absence of growth in *X*_2_ and a high switching rate from *X*_1_, the model anticipates no regime of stable spore concentration.

Experimental observations aligned with these predictions: at high dilution rates, residual glucose remained above the switching threshold and sporulation was not activated, yielding exclusively *X*_1_ biomass (**Fig. 2**). Reducing *D* caused residual glucose to fall below *K*_*I*_, initiating bursts of sporulation from *X*_1_ to *X*_2_. These oscillatory transitions persisted for at least five residence times before gradually decreasing in amplitude.

**Figure 2:**
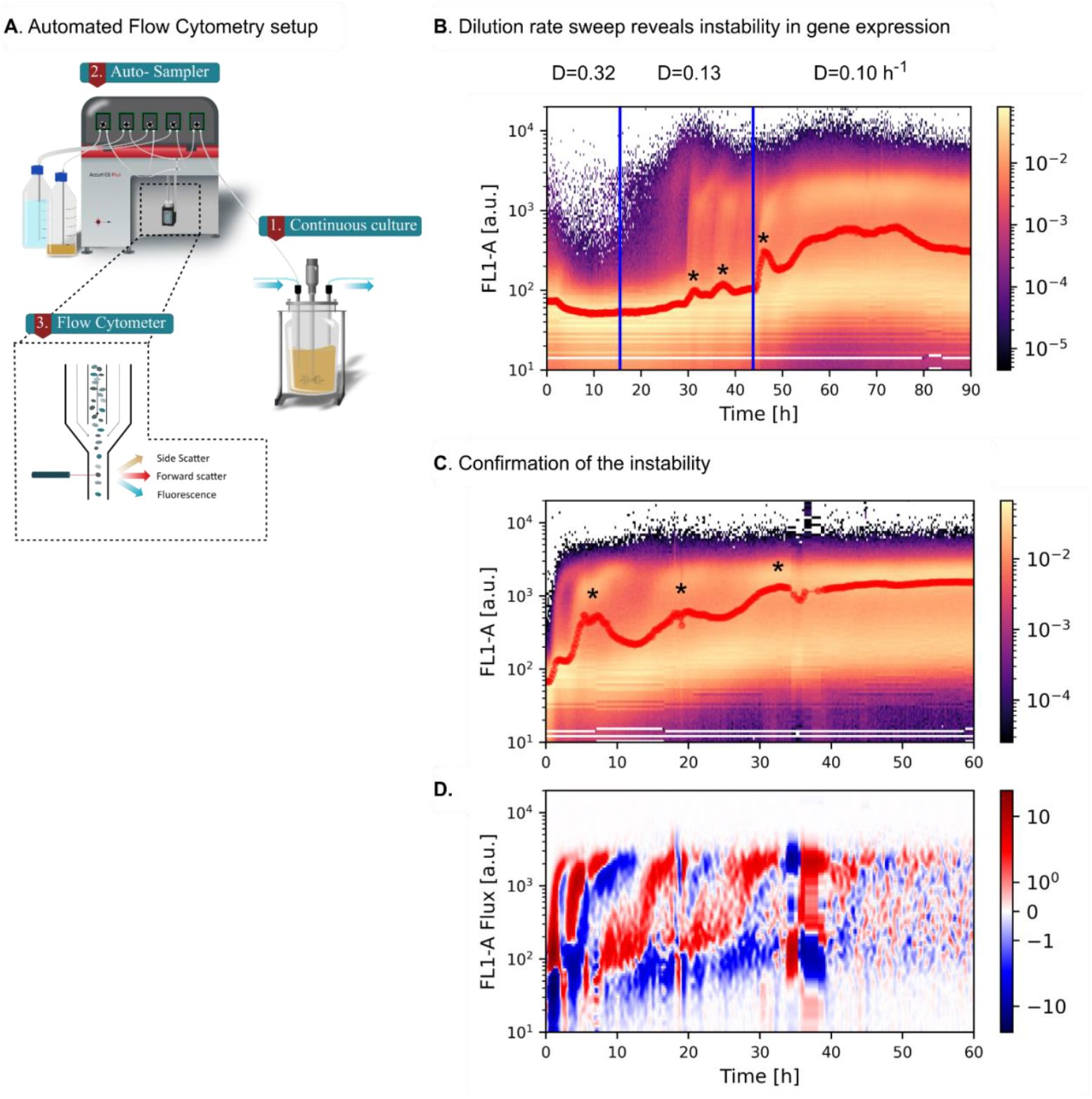
Mapping of cell population dynamics using automated flow cytometry reveals spontaneous oscillations in the activation of the *spoIIE* sysem during continuous cultivation of *B. subtilis*. **(A)** Schematic of the automated flow cytometry setup. Samples are withdrawn directly from the bioreactor and analyzed at single-cell resolution every 15 min. **(B)** Time–fluorescence density plots, where brighter colors indicate higher cell counts. The black line marks the 75th percentile of fluorescence; large oscillations are indicated by asterisks. In agreement with model predictions, gene expression dynamics vary with the dilution rate (*D*): at *D* = 0.32 h^−1^, cells remain vegetative, whereas lowering *D* to 0.13 h^−1^ triggers sporulation bursts detected by activation of the *P*_*spoIIE*_*::gfp* reporter. These bursts diminish in amplitude upon further reduction to *D* = 0.10 h^−1^. **(C)** Prolonged chemostat cultivation at *D* = 0.11 h^−1^ confirmed the persistence of oscillatory sporulation. **(D)** Differential fluorescence maps showing the increase (red) or decrease (blue) in cell counts per fluorescence bin between successive time points, smoothed for clarity. Until ∼40 h of continuous culture, clear waves of sporulation are observed. Color units are arbitrary.

## Discussion

Oscillatory behaviors in biology have long been associated with dedicated genetic oscillators, external forcing, or a combination of both^1 29 30^. From natural systems such as circadian clocks and cell cycles to synthetic designs like the repressilator^19^, the prevailing assumption has been that sustained oscillations require an explicit genetic architecture to generate them. Our work challenges this paradigm by demonstrating that oscillations can arise spontaneously in constant environments, purely from the interplay between cellular physiology and environmental feedback.

Our results demonstrate that even gene networks lacking intrinsic oscillatory architecture can generate sustained oscillations in the absence of external forcing. These oscillations emerge from the interplay of two independent but complementary mechanisms. The first is the impact of gene expression on cellular growth. This phenomenon is widespread: many natural circuits, such as sporulation in *B. subtilis*, and synthetic systems, such as burdensome protein overproduction^21^, can impose substantial growth penalties^22^. Similar effects are conceivable for genes affecting persistence traits, such as biofilm formation, where the “cost” to growth is offset by improved survival in specific environments^31^.

The second requirement is environmental feedback, in which cells not only sense but also modify the concentration of their inducers. This feedback occurs when cells consume, degrade, or produce the relevant inducer, thereby reshaping their own stimulus landscape. In our case study, sporulation reduced glucose uptake, allowing residual glucose to rise above the sporulation threshold and thereby suppressing further induction in non-stressed cells. A similar mechanism was reported for a T7 expression system under lactose control^31^, where growth inhibition from burdensome protein production increased residual glucose levels and suppressed induction via carbon catabolite repression.

Fundamentally, the oscillations observed here represent system-level circuits: emergent behaviors arising from population–environment interactions rather than from the regulatory topology of an isolated genetic network. Whereas genetic oscillators are defined by well-mapped gene–protein interactions^16 25^, system-level oscillators depend on how those interactions couple to resource flows and environmental feedback^32 31 33^. The example we present is relatively simple, but more complex interactions, especially where multiple gene networks intersect through shared environmental variables, could produce rich and unexpected dynamical patterns.

The emergence of oscillations in the absence of an explicit genetic oscillator has several important implications. First, it challenges the assumption, fundamental to chemostat theory^26 34^, that constant environmental conditions necessarily yield steady-state population structures and gene expression profiles. Second, it suggests that similar instabilities may occur in other long-term cultures whenever environmental feedback is tightly coupled to a phenotypic switch that significantly alters resource consumption. This principle could extend to diverse microbial systems, from industrial bioprocesses^35^ to natural microbial communities^36 37^, where environmental cues and physiological states are dynamically linked.

## Materials and Methods

### Mathematical Model Solution

The dynamic model describing growth and stress-induced phenotypic switching was solved analytically, as detailed in the Supplementary Information (**SI Appendix**, “Stress model in continuous culture”). To this end, we first considered separately the regimes with or without switching toward the stressed phenotype and, subsequently, merged both solutions to obtain a complete analytical description. The approach was then generalized to exponential fed-batch conditions. Additional analyses regarding the model’s domain of validity and biological assumptions are also provided in the SI.

The analytical solutions revealed that, for specific combinations of biological parameters (growth rates, switching kinetics) and operational conditions (dilution rate), steady-states do not exist, indicating transitions to oscillatory dynamics or population wash-out.

Model predictions were verified through numerical simulations of a continuous culture, performed in MATLAB using the delay differential equation solver *dde23*. Parameter values were selected based on experimental measurements obtained with our *Bacillus subtilis* strain, including a 30-min delay between the environmental stress threshold (substrate limitation) and activation or repression of the switch to the stressed phenotype.

### Experimental Design

The objective of this study was to elucidate how regular gene-expression oscillations can emerge in the absence of dedicated oscillator motifs or engineered external feedback. We used *Bacillus subtilis* sporulation as a model system for stress-induced phenotypic transitions. To parameterize the model, we first experimentally quantified growth and switching characteristics, as described in the “Growth and switching parameter determination” section of the SI. Using these parameters and the analytical framework described above, we identified dilution-rate regimes predicted to either suppress sporulation or induce sustained oscillations. Chemostat experiments were then performed at representative dilution rates within these regimes. Sporulation dynamics were monitored using our custom automated flow-cytometry setup, in combination with an engineered reporter strain producing GFP upon activation of the sporulation program. This setup enabled single-cell-resolved tracking of phenotypic transitions and direct validation of model predictions.

### Strain and Growth media

All experiments were performed in minimal mineral media containing (in g/l): K_2_HPO_4_: 14.6; NaH_2_PO_4_⋅2H_2_O: 3.6; Na_2_SO_4_: 2; (NH_4_)_2_SO_4_: 2.47; NH_4_Cl: 0.5; (NH_4_)2H-citrate: 1; glucose: 5, thiamine: 0.01, tryptophan: 0.05. The medium is supplemented with a trace element solution totaling 11 ml/l assembled from the following solutions (in g/l), 3/11 of FeCl_3_⋅6H_2_O: 16.7, 3/11 of EDTA: 20.1, 2/11 of MgSO_4_: 120 and 3/11 of a metallic trace element solution. The metallic trace element solution contains (in g/L): CaCl_2_⋅2H_2_O: 0.74; ZnSO_4_⋅7H_2_O: 0.18; MnSO_4_⋅H_2_O: 0.1; CuSO_4_⋅5H_2_O: 0.1, CoSO_4_⋅7H_2_O: 0.21. Both the trace element solution and the amino acids were filter sterilized (0.22 μm). To monitor the switching to the sporulating phenotype, *Bacillus subtilis* 168 was modified to express *gfpmut2* under the control of the SpoIIE promoter. The strain, *B. subtilis* 168 P_spoIIE:gfpmut2_, was built by AmyE chromosomal integration of a cassette containing the promoter, gfpmut2 and the selection maker, kanamycin.

### Cultivation Conditions

Precultures were started from single colonies picked from a LB plate and grown overnight. To avoid starting bioreactor cultivations with sporulating populations, the precultures were done in overfilled shake flasks (20 % of filled volume) without baffles to reach oxygen limitation before glucose. All bioreactor cultivations were performed at 37°C, pH 7 and aeration at 1 VVM. For growth and switching parameters determination, experiments were performed in DASBox mini - bioreactors (Eppendorf) with a stirring speed of 400 rpm and a cultivation volume of 160 ml. Continuous cultivations were performed in Bionet F1 bioreactor with a cultivation volume of 1L at an agitation speed of 1000 min^-1^. The transition from batch to continuous was triggered once dissolved oxygen rose, signaling the transition from exponential to stationary phase.

Bioreactor cultivations were monitored with a custom-made sampling device (the Segregostat) that automatically draws a sample from the reactor and dilutes it with PBS before flow cytometry (FC) analysis (BD Accuri, C6 for the parameter determination and C6+ for the continuous cultures). A total of 40,000 cells was analyzed in each sample where the FL1-A channel is used to visualize the GFP content.

### Statistical Analysis

The switching and growth parameters were determined in triplicate cultivations. The cultivations to test the impact of dilution rates were performed in duplicates and the dilution rate leading to instability was confirmed one last time. During all cultivations, samples were automatically taken every 15 minutes and 40,000 cells were analyzed.

## Acknowledgments

We thank Laurie Josselin, Juan Andres Martinez, Mélanie Grégoire and Mathéo Delvenne for commenting the article.

## Author contributions

V.V. contributed equally to this work with L.H. V.V. and L.H. developed the model, performed the experiments and drafted the manuscript. F.D. secured related fundings, wrote and reviewed the manuscript.

## Data and materials availability

The automated flow cytometry data and associated code have been deposited on https://zenodo.org/records/13803914). The data analysis code relies on the previously published toolbox available at https://gitlab.uliege.be/mipi/published-software/mbiomas-core.

