## Supplementary material for "Biological oscillations without genetic oscillator or external forcing"

bioRχiv
THE PREPRINT SERVER FOR BIOLOGY

### Supplementary materials for
### **Biological oscillations without genetic oscillator or external forcing**

Second version

Vincent Vandenbroucke[a,1], Lucas Henrion[a,1], and Frank Delvigne[a,2]

[1] V.V. and L.H. contributed equally to this work.

[a] Terra Research and Teaching Centre, Microbial Processes and Interactions (MiPI), Gembloux Agro-Bio Tech, University of Liège, Gembloux, Belgium

27th October 2025

**2 Limitations of the model and additional notes 11**

**3 Exponential fed-batch 12**

**4 Materials and Methods 13**

**6 Dataset legends 25**

**References 26**

### 1 Stress model in continuous culture

A chemostat is usually modelled with the following equations:

$$\frac{\mathrm{d}S}{\mathrm{d}t} = (S_{in} - S)D - \frac{\mu X}{Y} \tag{S.1}$$

$$\frac{\mathrm{d}X}{\mathrm{d}t} = (\mu - D)X \tag{S.2}$$

However, often, cells have a second state that can be induced when they are stressed, in which case their growth rate decreases. We can represent those two states in a chemostat with the following equations, if we consider that the stressed state is irreversible and cannot go back to non-stressed. As the time required to dilute the stress proteins is quite long, and made longer by the lower growth rate caused by stress, switching to a stress state is much faster and this is a reasonable assumption for a simplified approach. That is without considering the fact that, in a chemostat with non-stressed cells, the stressed-cells would be overtaken slowly and, if not renewed, are likely to simply be washed out before they can go back to their non-stressed state.

$$\frac{\mathrm{d}S}{\mathrm{d}t} = (S_{in} - S)D - \frac{\mu X_1}{Y_{X1}} - \frac{k\mu X_2}{Y_{X2}} \tag{S.3}$$

$$\frac{\mathrm{d}X_1}{\mathrm{d}t} = (\mu - D)X_1 - H_1(S)X_1 \tag{S.4}$$

$$\frac{\mathrm{d}X_2}{\mathrm{d}t} = (k\mu - D)X_2 + H_1(S)X_1 \tag{S.5}$$

$$\mu = \mu_{max}\frac{S}{K_S + S} \tag{S.6}$$

$$\mu_{max,1} = \mu_{max} \tag{S.7}$$

$$\mu_{max,2} = k\mu_{max} \tag{S.8}$$

$$H_1(S) = \begin{cases} H & \text{if } S \leq K_I \\ 0 & \text{if } S > K_I \end{cases} \tag{S.9}$$

Where the variables are defined as:

| | |
|---|---|
| $D$ | the dilution rate. |
| $k$ | the ratio of the growth rate of the slower phenotype to the growth rate of the faster phenotype. |
| $K_I$ | the threshold substrate concentration for the switch. |
| $K_S$ | the half-saturation constant of the Monod equation. |
| $H$ | the switching rate of the faster to the slower phenotype. |
| $H_1(S)$ | the switch function from the faster to the slower phenotype. |
| $\mu$ | the instantaneous growth rate of the faster phenotype. |
| $\mu_{max}$ | the maximum growth rate of the faster phenotype. |
| $S$ | the substrate (usually glucose) concentration in the reactor. |
| $S_{in}$ | the substrate concentration in the feed. |
| $t$ | the time. |
| $X_1$ | the concentration of the faster phenotype in the reactor. |
| $X_2$ | the concentration of the slower phenotype in the reactor. |
| $Y_{X1}$ | the yield of the faster phenotype. |
| $Y_{X2}$ | the yield of the slower phenotype. |

We can note the following conditions on the parameters based on the parameters:

- All variables are positive due to the constraints of reality and considering growing phenotypes.
- $D$, $Y_{Xi}$, $\mu_{max}$, $K_S$, $K_I$ and $H$ are strictly positive.

Considering those equations for a chemostat, the question that now occurs is:
What happens at steady state?

Due to the switch to a stress conditions, we have two kinds of steady-state to consider: ones where the substrate concentration is above the threshold $K_I$, and ones where the substrate concentration is below the threshold $K_I$. Separating those cases let us replace $H_1(S)$ by $H$ or 0 in the equations, greatly simplifying the system. Note that by definition, the steady-state requires the residual substrate concentration $S$ to be constant, therefore all the possible steady-state can be found by splitting the equations this way.

To jump to the solution of the steady-state, go to section 1.3.

#### 1.1 When $S > K_I$

In this case, at steady-state (derivatives are null), there is no switching and we can replace equations S.3 to S.5 by the following:

$$0 = (S_{in} - S)D - \frac{\mu X_1}{Y_{X1}} - \frac{k\mu X_2}{Y_{X2}} \tag{S.10}$$
$$0 = (\mu - D)X_1 \tag{S.11}$$
$$0 = (k\mu - D)X_2 \tag{S.12}$$

According to the latter 2 equations, there are 4 possible cases:

1. $X_1 = 0$ and $X_2 = 0$.
2. $X_1 = 0$ and $D = k\mu$.
3. $X_2 = 0$ and $D = \mu$.
4. If $k = 1$, $D = \mu$

These are the expected steady-states and show already known results: in the absence of switching, at steady-state, either the two populations are washed out or not initially present (case 1); and there is an unstable case where if both phenotypes have exactly equal growth rates, they can both survive, in any proportion as they behave as a single population and this whole model becomes irrelevant (case 4).

For the sake of this argument, we can consider $k \neq 1$ as that is a trivial and uninteresting case.

Both phenotypes cannot coexist in this case if $k \neq 1$, as one existing, in the absence of switching, imposes a specific dilution rate that implies, at steady-state, the other no being there.

The last case, a wash-out of both/either phenotype, will always happen for a population if $(\mu - D) = 0$ is impossible. To show when it is forced, finding out when it is possible is the easiest way.

Then, re-writing equations S.10 to S.12 for a steady-state with a single phenotype:

$$(S_{in} - S)D = \frac{X_i}{Y_{Xi}} \mu_{max,i} \frac{S}{K_S + S} \tag{S.13}$$
$$\mu_{max,i} \frac{S}{K_S + S} = D \tag{S.14}$$

where i indicates either phenotype, whichever is present since we established that in this case only one can be present.

Since $Y_{Xi}$, $D$ and $K_S$ are strictly positive, we can deduce:

$$\mu_{max,i} \frac{S}{K_S + S} = D \quad \Rightarrow \quad \frac{D}{\mu_{max,i}} = \frac{S}{K_S + S} \tag{S.15}$$
$$S_{in} - S = \frac{X_i}{Y_{Xi}} \quad \Rightarrow \quad S = S_{in} - \frac{X_i}{Y_{Xi}} \tag{S.16}$$

Therefore:

$$
\begin{aligned}
\frac{D}{\mu_{max,i}} &= \frac{S}{K_S + S} \\
&= 1 - \frac{K_S}{K_S + S} \\
\Leftrightarrow \qquad \frac{K_S}{K_S + S} &= 1 - \frac{D}{\mu_{max,i}} \\
&= \frac{\mu_{max,i} - D}{\mu_{max,i}} \\
\Leftrightarrow \qquad K_S + S &= \frac{\mu_{max,i} K_S}{\mu_{max,i} - D} \\
\Leftrightarrow \qquad S &= \frac{\mu_{max,i} K_S}{\mu_{max,i} - D} - K_S \\
&= \frac{D K_S}{\mu_{max,i} - D} \qquad \text{(S.17)}
\end{aligned}
$$

Plugging equation S.17 into equation S.16 and solving for $X_i$ gives:

$$
\begin{aligned}
S_{in} - \frac{D K_S}{\mu_{max,i} - D} &= \frac{X_i}{Y_{Xi}} \\
X_i &= Y_{Xi} \left( S_{in} - \frac{D K_S}{\mu_{max,i} - D} \right) \qquad \text{(S.18)}
\end{aligned}
$$

Thus, eventually, from equations S.15, S.16, S.17, and S.18 we get:

$$
\begin{cases}
D &= \mu_{max,i} \frac{S}{K_S + S} \\
S &= \frac{D K_S}{\mu_{max,i} - D} = S_{in} - \frac{X_i}{Y_{Xi}} \\
X_i &= Y_{Xi} \left( S_{in} - \frac{D K_S}{\mu_{max,i} - D} \right) = Y_{Xi}(S_{in} - S)
\end{cases} \qquad \text{(S.19)}
$$

at steady states, for cases 2 and 3. And since $S \geq 0$ and $X_i \geq 0$, for this to be possible, we need:

- $\mu_{max,i} > D$ for $S \geq 0$. When a given phenotype is initially present, a wash-out and no biomass is leftover at steady-state (it becomes case 1) if the maximum growth rate of the phenotype is lower than the dilution rate (as expected).
- $S_{in} > {}^{DK_S}/_{(\mu_{max,i} - D)}$ for $X_i > 0$. This can be rearranged (as per standard chemostat equations) as $D < \mu_{max,i} {}^{S_{in}}/_{(K_S + S_{in})} < \mu_{max,i}$, dilution rate above which there is no glucose consumption at equilibrium as there is not enough substrate to sustain the population.

Those equations and deductions are only valid when $S > K_I$, or:

$$
\frac{D K_S}{\mu_{max,i} - D} > K_I \qquad \text{(S.20)}
$$

Firstly, we established that, when there is no wash-out, and there is biomass at steady-state, $\mu_{max,i} > D$ is required. Therefore, the left hand-side of the inequality is always positive.

We can examine when transform the inequality to examine when it is possible or:

$$
\begin{aligned}
& \frac{DK_S}{\mu_{max,i} - D} > K_I \\
\Leftrightarrow \quad & DK_S > K_I(\mu_{max,i} - D) && \text{since: } \mu_{max,i} > D \\
\Leftrightarrow \quad & DK_S + DK_I > K_I\mu_{max,i} \\
\Leftrightarrow \quad & D(K_S + K_I) > K_I\mu_{max,i} \\
\Leftrightarrow \quad & D > \frac{K_I\mu_{max,i}}{K_S + K_I} && \text{since: } K_S, K_I > 0
\end{aligned}
\tag{S.21}
$$

Therefore, steady-state without switching can be reached when $\mu_{max,i}{}^{S_{in}}/(S_{in} + K_S) > D > \mu_{max,i}{}^{K_I}/(K_S + K_I)$ for each phenotype if the three parameters allow (although, for the stressed phenotype, we are assuming that there is no switch back to the first one, which is probably wrong if waiting long enough in this high substrate concentration case).

As a summary of this section, we have for each case described at the start:

1. Wash out of all the cells initially present, when $D > \mu_{max,i}\frac{S_{in}}{(S_{in}+K_S)}$ for all $i$ phenotypes non-zero in the initial conditions.

2. a steady state with only the second phenotype present, which could happen if all cells switched to the second phenotype, and never switched back, maybe in the case of differentiation:

$$
\begin{cases}
X_1 = 0 \\
D = k\mu_{max}\ {}^{S}/K_S + S \\
S = {}^{DK_S}/k\mu_{max} - D = S_{in} - {}^{X_2}/Y_{X2} \\
X_2 = Y_{X2}(S_{in} - S) \\
0 \le S \le S_{in} \\
k\mu_{max}{}^{S_{in}}/(S_{in} + K_S) > D > k\mu_{max}{}^{K_I}/(K_S + K_I)
\end{cases}
\tag{S.22}
$$

3. A steady state with only the first phenotype present:

$$
\begin{cases}
X_2 = 0 \\
D = \mu_{max}\ {}^{S}/K_S + S \\
S = {}^{DK_S}/(\mu_{max} - D) = S_{in} - {}^{X_1}/Y_{X1} \\
X_1 = Y_{X1}(S_{in} - S) \\
0 \le S \le S_{in} \\
\mu_{max}{}^{S_{in}}/(S_{in} + K_S) > D > \mu_{max}{}^{K_I}/(K_S + K_I)
\end{cases}
\tag{S.23}
$$

#### 1.2 When $S \leq K_I$

A similar reasoning can be applied to the alternative case, where the switching always occurs.

At steady-state, we get:

$$0 = (S_{in} - S)D - \frac{\mu X_1}{Y_{X1}} - \frac{k\mu X_2}{Y_{X2}} \tag{S.24}$$
$$0 = (\mu - D - H)X_1 \tag{S.25}$$
$$0 = (k\mu - D)X_2 + HX_1 \tag{S.26}$$

The equations can be solved easily when $X_1 = 0$ and $X_2 = 0$, as this means there is no biomass. Notably, this case is incompatible with the assumption that there is switching regardless as $S_{in} = S$ in that case, and in a bioreactor $S_{in} > K_I$, so we come back to the previous section.

In case $X_1 = 0$, the equations become identical to the previous section. The only difference is the condition of existence derived at the end, which becomes $D < \mu_{max,i}{}^{K_I}/_{K_S + K_I}$ instead. This completes all cases where only the second phenotype is present with a region where it is actually likely that all cells stay in the second phenotype, as the substrate concentration would be too low to permit switching back to the first phenotype.

Thus the only case of interest to discuss here is when both $X_1$ and $X_2$ are non-zero.

Re-arranging equations S.24 to S.26 with those assumptions:

$$(S_{in} - S)D = \frac{\mu X_1}{Y_{X1}} - \frac{k\mu X_2}{Y_{X2}} \tag{S.27}$$
$$\mu = \mu_{max}\frac{S}{K_S + S} = D + H \tag{S.28}$$
$$(D - k\mu)X_2 = HX_1 \tag{S.29}$$

Interestingly, equation S.28 implies that the first phenotype, through $D$, $H$, $\mu_{max}$ and $K_S$, fully determines the substrate concentration, with only $H$ differing from a normal chemostat and no direct influence of the phenotype ratio in the culture.

Based on equation S.28, we can deduce:

$$S = \frac{(K_S + S)(D + H)}{\mu_{max}}$$
$$\Leftrightarrow \quad S\left(1 - \frac{D + H}{\mu_{max}}\right) = K_S\frac{D + H}{\mu_{max}}$$
$$S\,\frac{\mu_{max} - D - H}{\mu_{max}} =$$
$$\Leftrightarrow \quad S = \frac{K_S(D + H)}{\mu_{max} - D - H} \tag{S.30}$$

Equation S.29 can easily be re-arranged into:

$$X_1 = \frac{D - k\mu}{H} X_2 \Leftrightarrow X_2 = \frac{H}{D - k\mu} X_1 \tag{S.31}$$

From there, we can plug into the remaining equation S.27:

$$\begin{aligned} & (S_{in} - S)\ D = \frac{\mu X_1}{Y_{X1}} + \frac{k\mu}{Y_{X2}}\ \frac{H}{D - k\mu}\ X_2 \\ \Leftrightarrow \quad & X_1 = \frac{(S_{in} - S)\ D}{\frac{\mu}{Y_{X1}} + \frac{k\mu H}{Y_{X2}\ (D - \mu k)}} \quad \text{(S.32)} \\ \text{and} \quad & (S_{in} - S)\ D = \frac{(D - k\mu)\mu}{HY_{X1}}\ X_2 + \frac{k\mu X_2}{Y_{X2}} \\ \Leftrightarrow \quad & X_2 = \frac{(S_{in} - S)\ D}{\mu\ \left(\frac{(D - k\mu)}{HY_{X1}} + \frac{k}{Y_{X2}}\right)} \quad \text{(S.33)} \end{aligned}$$

We can thus write the solution for this case of the steady-state as:

$$\begin{cases} S & = \frac{K_S(D+H)}{\mu_{max} - D - H} \\ \mu & = \mu_{max} \frac{S}{K_S + S}\ =\ D + H \\ X_1 & = \frac{(S_{in} - S)\ D}{\mu/Y_{X1} + k\mu H/Y_{X2}\ (D - \mu k)} \\ X_2 & = \frac{(S_{in} - S)\ D}{\mu\ ((D - k\mu)/(HY_{X1}) + k/Y_{X2})} \\ \frac{X_1}{X_2} & = \frac{D - k\mu}{H} \end{cases} \tag{S.34}$$

For this solution to be possible, we need:

- $S > 0$
- $X_1 > 0$
- $X_2 > 0$

Since $X_1 = X_2 {}^{(D - k\mu)}/_{H}$, we need for both to be positive:

$$D > k\mu \tag{S.35}$$

Since $D + H = \mu$, we actually need:

$$\frac{D}{D + H} > k \tag{S.36}$$

Note that this condition can only be true if $k < 1$, as the left hand side is always smaller than 1. Thus, if the induced phenotype were to grow faster than the initial one, it would be impossible for the two phenotypes to coexist in the reactor. In this case, we are considering a stress phenotype, so $k < 1$ is a reasonable assumption.

In other musings, this condition means at its origin that the dilution rate will be faster than the instantaneous growth rate of the second phenotype.

The condition can be easily re-arranged in terms of $D$ as well:

$$\frac{D}{D+H} > k \qquad \Leftrightarrow \qquad D > \frac{kH}{1-k} \tag{S.37}$$

This ensures that both $X_1$ and $X_2$ have the same sign. To ensure they are positive, we need:

$$S_{in} > S \tag{S.38}$$

which is always true if cells grow.

Finally, we need for $S \geq 0$:

$$D < \mu_{max} - H \tag{S.39}$$

The latter also implies that, for both phenotypes to coexist,

$$\mu_{max} > H \tag{S.40}$$

is required.

Those conditions, combined with the one required by this section, $S \leq K_I$, are the only conditions required for the existence of a steady-state with both phenotypes present.

The latter condition can be developed:

$$\begin{aligned}
& \frac{K_S(D+H)}{\mu_{max} - (D+H)} < K_I \\
\Leftrightarrow \quad & K_S(D+H) < K_I\ (\mu_{max} - (D+H)) \qquad \text{"<" sign unchanged since (S.39)} \\
\Leftrightarrow \quad & DK_S + HK_S < \mu_{max}K_I - DK_I - HK_I \\
\Leftrightarrow \quad & D(K_S + K_I) < \mu_{max}K_I - H(K_S + K_I) \\
\Leftrightarrow \quad & D < \frac{\mu_{max}K_I}{K_S + K_I} - H
\end{aligned} \tag{S.41}$$

Similarly to condition S.40, we can deduce that

$$\mu_{max}\frac{K_I}{K_S + K_I} > H \tag{S.42}$$

from the previous condition.

Note that condition S.41 is stronger than condition S.39, as:

$$\begin{aligned} \mu_{max} - H &> \frac{\mu_{max} K_I}{K_S + K_I} - H \\ \Leftrightarrow \qquad \mu_{max} &> \mu_{max} \frac{K_I}{K_S + K_I} \end{aligned} \tag{S.43}$$

Inequation S.43 is always true, as $\frac{K_I}{(K_S+K_I)}$ is smaller than one, so condition S.39 is irrelevant. The deduced condition on the relationship is similiarly stronger in (S.42) than in (S.40).

In summary, the steady-state with both phenotypes present is possible with the non-trivial conditions:

- ${}^{(\mu_{max} K_I)}\!/\!_{(K_S+K_I)} - H > D > {}^{kH}\!/\!_{(1-k)}$
- $\mu_{max} {}^{K_I}\!/\!_{(K_S+K_I)} > H$

#### 1.3 Solution of the steady-state

Overall, the steady-state of the cells can be described as a function of an increasing dilution rate:

$$\begin{array}{rll} X_2 \text{ only}: & \begin{cases} D < k\mu_{max} {}^{K_I}\!/\!_{(K_S+K_I)} \\ k\mu_{max} {}^{K_I}\!/\!_{(K_S+K_I)} < D < k\mu_{max} {}^{S_{in}}\!/\!_{(K_S+S_{in})} \\ k\mu_{max} {}^{S_{in}}\!/\!_{(K_S+S_{in})} < D \end{cases} & \begin{array}{l} \text{Only } X_2 \text{ is present, no induction} \\ \text{Only } X_2 \text{ is present, but } S > K_I \\ \text{Wash-out of } X_2 \text{ only} \end{array} \\ X_1 \text{ and } X_2: & \left\{ {}^{kH}\!/\!_{(1-k)} < D < \mu_{max} {}^{K_I}\!/\!_{(K_S+K_I)} - H \right. & {}^{X_1}\!/\!_{X_2} = {}^{(D-k\mu)}\!/\!_{H} \\ X_1 \text{ only}: & \begin{cases} \mu_{max} {}^{K_I}\!/\!_{(K_S+K_I)} < D < \mu_{max} {}^{S_{in}}\!/\!_{(K_S+S_{in})} \\ \mu_{max} {}^{S_{in}}\!/\!_{(K_S+S_{in})} < D \end{cases} & \begin{array}{l} \text{Only } X_1\text{, no induction} \\ \text{Wash-out} \end{array} \end{array}$$

For $X_2$ only, while there are multiple cases, when those cases are overlapping with another region, then that other region is most likely to be reached instead: other regions include $X_1$, and while our assumptions include that there is no switch from $X_2$ to $X_1$, that assumption would likely prove wrong in the long term. In particular, the region where $X_2$ is the only phenotype present and $S > K_I$ (inhibiting the switch from $X_2$ to $X_1$) is probably not stable and thus we will not consider it further.

Interestingly, since no assumption can be made on the relative values of the different parameters, the different regions do not always define the entire range of dilution rates.

Between the regions where only $X_2$ and both phenotypes are present, there may be a region without steady-state if ${}^{kH}\!/\!_{(1-k)} > k\mu_{max} {}^{K_I}\!/\!_{(K_S+K_I)}$. Between those two thresholds of dilution rates, the substrate concentration would be too high to keep inducing the second phenotype.

Between the regions where only $X_1$ and both phenotypes are present, there may be a region without steady-state if $\mu_{max} {}^{K_I}\!/\!_{(K_S+K_I)} > \max\{\mu_{max} {}^{K_I}\!/\!_{(K_S+K_I)} - H \ , \ k \cdot (\mu_{max} {}^{K_I}\!/\!_{(K_S+K_I)})\}$, where the second term of the maximum is in case the $X_2$ only region reaches beyond the cohabitation region. Whichever is the maximum, this inequality is always true (because of the $-H$ or because $k < 1$). Therefore, there is always a region without steady-state when decreasing the dilution rate and the stressed phenotype starts to be induced induced. This interval is the one discussed in the main paper.

In those regions without steady-state, any attempt to reach steady-state when $S > K_I$ causes the biomass to increase and $S$ to decrease. When $S < K_I$, the cells start switching and substrate consumption decreases, causing $S$ to increase above $K_I$ again. Since cells do not respond instantaneously to change, and in this region where $S$ is close to $K_I$ small changes in concentration can cause large changes in responses, the cells may oscillate between the two states.

### 2 Limitations of the model and additional notes

There are a few additional things that can be added on this model, in no particular order:

- While it was derived for a chemostat, it can be applied to an exponential fed-batch reactor as well (see section 3). For a fed-batch with less than exponential feeding, the effective growth rate decreases over time. In other words, it would not be a question of whether the region without equilibrium is reached, but when. Then the second question is whether that will cause oscillations or not, which will depend on the exact biological system, and how long the cells stay in the problematic region.
- This model predicts that there is a region where there is no steady-state when there are sharp transitions in cell metabolism, such as with stress induction, or induction of burdensome production with lactose. Oscillations may appear thanks to the delay in response time of the cells, and very small changes in substrate concentration causing large changes in cell response. However, the switch is not a perfect step function, and the delay will be shorter or longer depending on the cells. Further, a constant switching speed may not be a good approximation, at least when close to the switching threshold. All in all, oscillations may appear, but they may be dampened, or the the system may stabilize quickly in an intermediate state that this model is incapable of representing.
- In the demonstration, we considered what happens when $K_I < S$ and $K_I > S$. When $K_I = S$, the details of what happen in the model depend on how $H_1(S)$ was defined (here, $H_1(K_I) = H$, therefore the $K_I = S$ would be the limit of the $K_I < S$ case). In practice, however, reality is not completely discontinuous, and this is the edge of an undefined region regardless, so the mathematics of this particular fictional point do not really matter, only the limits of the region between cases does.
- We consider here that both phenotypes are growing, no matter how slowly. If the slower phenotype were to have a death rate instead ($k < 0$), several things would happen:
  - No region would exist where only the second phenotype can survive, as even a batch culture would eventually have no cells left. This is expressed by condition S.37 being always true.
  - Nothing else would change, as in no place in section 1.2 is there any division by $k$ for any condition. We can therefore expect exactly the same dynamics and region of uncertainty as with $k \geq 0$.

- For a case such as the induction of burden with a second carbon source (such as the induction of BL21 with lactose (co-feeding of glucose and lactose), where oscillations were observed as well [1]), considering co-consumption of carbon sources, the inequations are inverted such that equations S.19 is valid only when $D < \mu_{max}{}^{K_I}/_{K_S + K_I}$, and equations S.34 is valid only when $\mu_{max} - H > D > \mu_{max}{}^{K_I}/_{K_S + K_I} - H$ and $D > H^{k}/_{(1-k)}$. Further, the case of only the second phenotype being present should be stable between $k\mu_{max}{}^{K_I}/_{(K_S + K_I)}$ and $k\mu_{max}$. Therefore, a steady-state (or more than one) always exists at least as long as the induced phenotype is not washed out, and oscillations may only be an intermediate state that will eventually stabilize. Accounting for those oscillations would require investigation of the exact dynamics of the system of interest, and probably require taking into account carbon catabolite repression (When induced, cells stop growing, this increases both glucose and lactose concentrations. Increasing glucose concentration would inhibit induction by carbon catabolite repression, from which similar dynamics to the stress case would follow).

### 3 Exponential fed-batch

The traditional equations for a fed-batch reactor are:

$$\frac{\mathrm{d}(SV)}{\mathrm{d}t} = F(t)\ S_{in} - \frac{\mu XV}{Y} \tag{S.44}$$

$$\frac{\mathrm{d}(XV)}{\mathrm{d}t} = \mu XV \tag{S.45}$$

$$\frac{\mathrm{d}V}{\mathrm{d}t} = F(t) \tag{S.46}$$

Where $F(t)$ is the flow rate added to the reactor, and $D$ is the expected growth rate of the cells. In case of exponential feeding, $V$ and $^{\mathrm{d}V}/_{\mathrm{d}t}$ are defined as:

$$V(t) = C_0 + C_1 e^{Dt} \tag{S.47}$$

$$\frac{\mathrm{d}V}{\mathrm{d}t} = C_1 D e^{Dt} \tag{S.48}$$

Where $C_0$ is the integration constant, so that $V(0) = V_0$ the initial volume, and $C_1$ modulates the state of the exponential. Traditionally, $C_1$ is defined as $V_0$ (see [2]), which makes $C_0 = 0$ and simplifies the equations significantly. In Öztürk, Çalık, and Özdamar [3], however, the authors report a common value of $C_1 = {}^{V_0 X_0}/_{S_{in} Y}$ for *B. subtilis*. We can note that, if the initial glucose concentration is equal to $S_{in}$, then we can approximate $C_1 \approx V_0$. Regardless of the exactitude of that approximation, for the fed-batch to make sense, we need $V_0 > C_0 > 0$. Therefore, since we are here interested in steady-states only, we can $t_0$ the original time of the reactor such that $V(t_0) = V_0$, which means that $C_0$ can arbitrarily set to 0 and the steady-states will be identical.

The equations for the fed-batch can be adapted like for the chemostat:

$$\frac{\mathrm{d}(SV)}{\mathrm{d}t} = F(t)\ S_{in} - \frac{\mu X_1 V}{Y_{X1}} - \frac{k\mu X_2 V}{Y_{X2}} \tag{S.49}$$

$$\frac{\mathrm{d}(X_1 V)}{\mathrm{d}t} = V X_1 \mu - H_1(S) X_1 V \tag{S.50}$$

$$\frac{\mathrm{d}(X_2 V)}{\mathrm{d}t} = V X_2 k\mu + H_1(S) X_1 V \tag{S.51}$$

$$\frac{\mathrm{d}V}{\mathrm{d}t} = F(t) = C_1 D e^{Dt} \tag{S.52}$$

$$V(t) = C_1 e^{Dt} \tag{S.53}$$

Those equations can be set back into concentrations by using the chain rule for $A = S, X_1, X_2$ in the following equation:

$$\frac{\mathrm{d}(AV)}{\mathrm{d}t} = \frac{\mathrm{d}A}{\mathrm{d}t} V + A \frac{\mathrm{d}V}{\mathrm{d}t} = V \frac{\mathrm{d}A}{\mathrm{d}t} + AF(t) \tag{S.54}$$

The equations become:

$$\frac{\mathrm{d}S}{\mathrm{d}t} = \frac{F(t)}{V}(S_{in} - S) - \frac{\mu X_1}{Y_{X1}} - \frac{k\mu X_2}{Y_{X2}} \tag{S.55}$$

$$\frac{\mathrm{d}X_1}{\mathrm{d}t} = (\mu - \frac{F(t)}{V} - H_1(S)) X_1 \tag{S.56}$$

$$\frac{\mathrm{d}X_2}{\mathrm{d}t} = (k\mu - \frac{F(t)}{V}) X_2 + H_1(S) X_1 \tag{S.57}$$

Since ${}^{F(t)}/_{V} = D$, these equation become exactly identical to a chemostat, and the same conclusions can be drawn.

### 4 Materials and Methods

#### 4.1 Strain and growth media

All experiments were performed in minimal mineral media containing (in g/l): $K_2HPO_4$: 14.6; $NaH_2PO_4 \cdot 2H_2O$: 3.6; $Na_2SO_4$: 2; $(NH_4)_2SO_4$: 2.47; $NH_4Cl$: 0.5; $(NH_4)_2H - \mathrm{citrate}$: 1; glucose: 5, thiamine: 0.01, tryptophan: 0.05. The medium is supplemented with a trace element solution totaling 11 ml/l assembled from the following solutions (in g/l), 3/11 of $FeCl_3 \cdot 6H_2O$: 16.7 , 3/11 of EDTA: 20.1, 2/11 of $MgSO_4$: 120 and 3/11 of a metallic trace element solution. The metallic trace element solution contains (in g/L): $CaCl_2 \cdot 2H_2O$: 0.74; $ZnSO_4 \cdot 7H_2O$: 0.18; $MnSO_4 \cdot H_2O$: 0.1; $CuSO_4 \cdot 5H_2O$: 0.1, $CoSO_4 \cdot 7H_2O$: 0.21. Both the trace element solution and the amino acids were filter sterilized (0.22 $\mu m$). To monitor the switching to the sporulating phenotype, *Bacillus subtilis* 168 was modified to express *gfpmut2* under the control of the *SpoIIE* promoter. The strain, *B. subtilis* 168 $P_{spoIIE} : gfpmut2$, was built by *AmyE* chromosomal integration of a casette containing the promoter, $gfpmut2$ and the selection maker, kanamycin.

#### 4.2 Cultivation conditions and flow cytometry analysis

Precultures were started from single colonies picked from a LB plate and grown overnight. To avoid starting bioreactor cultivations with sporulating populations, the precultures were done in overfilled shake flasks (20 % of filled volume) without baffles to reach oxygen limitation before glucose. All bioreactor cultivations were performed at 37°c, pH 7 and aeration at 1 VVM. For growth and switching parameters determination, experiments were performed in DASBox mini - bioreactors (Eppendorf) with a stirring speed of 400 rpm and a cultivation volume of 160 ml. Continuous cultivations were performed in Bionet F1 bioreactor with a cultivation volume of 1 l at an agitation speed of 1000 rpm. The transition from batch to continuous was triggered once oxygen rose, signaling the transition from exponential to stationary phase.

Bioreactor cultivations were monitored with a custom made sampling device (the Segregostat) that automatically draws a sample from the reactor and dilutes it with PBS before flow cytometry (FC) analysis (BD Accuri, C6 for the parameter determination and C6+ for the continuous cultures). A total of 40,000 cells were analyzed in each sample where the FL1-A channel is used to visualize the GFP content.

#### 4.3 Growth and switching parameter determination

A triplicate batch cultivation in the DASBox was performed to estimate $\mu_{max}$ and switching ($K_I$, $H$) parameters of our strain. To this end, each bioreactor was inoculated with a OD of roughly 0.08 and samples were collected throughout the batch to assess the OD and the residual glucose concentration. The residual glucose concentration was determined using a D-glucose enzymatic assay kit from Megazyme in 98-well plates with a plate reader. During the batch, FC analysis was performed with our custom platform, the Segregostat, to observe when and at what speed cells were sporulating. The $K_S$ was defined based on the analysis the residual glucose concentration at steady state at a high dilution rates (where cells are not sporulating) during the chemostats with varying dilution rates.

The following parameters were measured or estimated:

- $H = 1.1$ h$^{-1}$ → Average from 0.83, 1.2, and 1.3 with r$_2$ = 0.98, 0.99 and 0.99 respectively (computed by linear regression over the time there is switching)
- $K_S = 0.14$ g/L → Based on the concentration at D=0.32 h$^{-1}$, measured as 0.2761 and 0.2432 g/L. From $D = \mu_{max} {}^{S}/_{(K_S+S)}$, we get $K_S = \mu_{max} {}^{S}/_{D} - S$
- $K_I \in [0.01, 0.1]$ g/L → Explained below
- $\mu_{max} = 0.49$ h$^{-1}$ → Average from 0.52, 0.47 and 0.48 with r$^2$ > 0.99 for all three (computed by linear regression over the time of exponential growth, see code)
- $k = 0$ → Sporulating cells are considered non-growing

To determine the start of sporulation, we considered the time at which the measured optical density dropped. That drop likely corresponds to a change in morphology of the cells. For that stretch of time, samples were taken more often than the automated flow cytometry, providing a better time resolution, in addition to having no absolutely no cross-contamination (using the segregostat with our DasGip in triplicate setup, a few percent leftover from each sample may be measured with the next sample), which is relevant when looking at very low percentages of induction. Based on these OD measurements, we considered the induction threshold was passed during the preceding measured time interval. This corresponds to 0.013 to 0.097 g/L, 0.009 to 0.068 g/L and 0.010 to 0.282 g/L for reactors 1, 2 and 3 respectively. Thus we estimated $K_I$ to be in the range of 0.01 to 0.1 g/L, although considering the cells need time to switch, $K_I$'s order of magnitude is likely the higher end of the range.

Based on these measurements, we could compute the unstable range of dilution rates for *B. subtilis*. The lower end of the range is at $D = 0$ $^{-1}$, as $k = 0$ and $(\mu_{max}K_I)/(K_S + K_I) - H < 0$ ($\mu_{max} < H$). The upper end depends on $K_I$. Based on the estimated range of $K_I$, it should be between $(\mu_{max}K_I)/(K_S + K_I) \in [0.03, 0.2]$ h$^{-1}$. The first replicate of the experiment showcasing a range of dilution rates showed the start of the instability at $D = 0.17$ h$^{-1}$, which would correspond to a $K_I$ of 0.075 g/L.

### 5 Code and Data Analysis

Three different codes were used for this paper, all are available in the supplementary dataset. First, a short Matlab code was used to simulate the system with a delay.

Second, a data analysis Python script was used to estimate the parameters from the experimental data. Due to its length, only the essential part of the code is presented in this document (the complete code is available in the supplementary dataset).

Third, a small data analysis Python script was used to plot the experimental data. The Python data analysis relies on the previously published toolbox [4] at `https://gitlab.uliege.be/mipi/published-software/mbiomas-core`. All the datasets used in this paper are available (`https://doi.org/10.5281/zenodo.13803913`).

#### 5.1 Matlab code

```
% system:
% dS= (Sin - S)D - muX1/YX1 - kmuX2/YX2
% dX1 = (mu - D)X1 - H1 (S)X1
% dX2 = (kmu - D)X2 + H1 (S)X1
% mu = mumax*S/(KS + S)
% mumax,1 = mumax
% mumax,2 = kmumax
%  if S <= KI
% H1 (S) = H
%  if S > KI
% H1 (S) = 0
```

```

sol = dde23(@diff_test1,[0.5],[0.02,2,0],[0,80])

% plot with X1 and X2 on left axis and S on right axis:
plot(sol.x,sol.y(2:3,:))
yyaxis right
plot(sol.x,sol.y(1,:),'k')
% add horizontal line at KI
hold on
plot([0,60],[0.075,0.075],'k--')
legend('X1','X2','S')

% save sol.x and sol.y to csv
writematrix([sol.x',sol.y'], 'test1.csv')

function diff=diff_test1(~, y, Z)
  % time unit: hour
  H=1.1;
  % k=0 so things easier
  KS=0.14;
  KI=0.075; % with D threshold=0.17 h-1 as experimented

  Sin=5;
  YX1=0.5; % standard value

  D=0.1; % dilution rate
  mu_max=0.49;

  diff=zeros(3,1);
  diff(1)= (Sin - y(1))*D-mumax*y(1)/(KS+y(1))*y(2)/YX1;
  diff(2)= (mumax*y(1)/(KS+y(1))-D)*y(2)-H*(Z(1)<KI)*y(2);
  diff(3)= H*(Z(1)<KI) * y(2) - D*y(3);
end
```

#### 5.2 Python script for data representation

```
import os
import datetime
import glob
import numpy as np
import pandas as pd
import mbiomas.pipes
import mbiomas.read
import mbiomas.fca
import matplotlib.pyplot as plt
```

```
from scipy.stats import linregress

mbiomas.pipes.write_version()

# standard reading data from dasgip:

def read_old_dasgip(folder, zero_time, time_zone, reactor_equivalence,
                    fields='all',
                    meta_encoding='utf-8',
                    time_offset_das=0):
    """
    Read data from the old DASGIP bioreactors (300 mL vessels from before 2020)
    This may include:
    * Online Flow cytometry data
    * Oxygen data, either in the root folder, or in an 'Oxygen' subfolder
        expected naming scheme:
        '*-chx.txt' where x is the probe number
    * Any number of offline data files, all the csv files.
        To indicate reactor number, the file should be named as:
        '*_x.csv' where n is the reactor number.
        (offline data is expected in local time)
        Metadata for those files should be named as the file, but with the
        extension `.meta.csv`.

    Parameters
    ----------
    folder : str
        Path to the folder containing the data files
    zero_time : datetime.Datetime or string
        Time at which the zero should be set. It is expected in local time.
    time_zone : str
        A number between -12 and 12, which indicates the
        difference between the zero time and the local time (UTC+2 → 2).
    reactor_equivalence : np.array
        Array with the equivalence between the different reactor numbers.
        Each line is a vessel.
        * The first column is the reactor number in the DASGIP software
        * The second column is the reactor number in the Oxygen software
        * The third column is the reactor number in the flow cytometry data
        * The fourth column is the reactor number in the offline data
        A '-1' indicates that the reactor is not present in the data.
        The order used is the one of the vessels in the matrix,
            top to bottom.
    fields : list
```

```
        List of fields to be read from the offline files. Will need to change so
        the default is all of them.
    meta_encoding : str
        Encoding of the offline data metadata files. Default is 'utf-8'
    time_offset_das : float
        Time offset in hours in the DasGIP computer. Default is 0. For example:
        Real time is 10:40, but the DasGIP computer is set to 10:50. Then the
        offset is 5/6-4/6=1/6=0.1667.
    """
    (... see supplementary dataset)

t=read_old_dasgip('batch',
                '2024-03-06 09:55',
                2,
                np.array(
                    [[-1,2,1,1],
                        [-1,3,2,2],
                        [-1,4,3,3]]),
                time_offset_das=-52/60)#8 minutes offset + 1h to real time

# remove useless columns
fields=['FL2-A', 'FL3-A', 'FL4-A', 'FL2-H', 'FL3-H', 'FL4-H', 'Width', 'Time']
for set_data in t:
    # for field in ['FL2-A', 'FL3-A', 'FL4-A',
    #                  'FL2-H', 'FL3-H', 'FL4-H', 'Width', 'Time']:
    set_data[3].remove_fields(fields)

# H time threshold (see processing later):
H_time_threshold=np.array([[12.6, 17],
                            [14.5, 18],
                            [14.2, 18]])

# fluorescence threshold
threshold=400

# save the measured parameters to a csv file through a pandas dataframe
parameters=pd.DataFrame(index=[0,1,2])

for i,set_data in enumerate(t):

    set_data[2], set_data[3],_=mbiomas.fca.data.clean(set_data[2], set_data[3])

    fig, ax=plt.subplots()

```

```
    plot=mbiomas.fca.plot.Plot(set_data[2])

    plot.time_scatter(ax,set_data[2],set_data[3],y_label='FL1-A',
        y_log_lim=[0,5])

    ## plot the percentage of cells above fluorescence threshold to determine
    ## H the switching rate

    fig, ax=plt.subplots()

    # make an empty DF to store the results
    percent_positive=pd.DataFrame(index=set_data[2].data["Time"])

    # add absolute time too
    # do it with a loop because it creates NaT otherwise
    # probably due to differing index formats
    # percent_positive.loc[:,'time']=set_data[2].data["Time"]
    # print(set_data[2].data["btimdilution"])
    # print(percent_positive)
    for j in range(len(percent_positive)):
        percent_positive.loc[percent_positive.index[j], 'Absolute time']= \
            set_data[2].data["btimdilution"][j]

    for j in range(len(percent_positive)):
        percent_positive.loc[percent_positive.index[j], 'percent']= \
                (set_data[3].data[j]['FL1-A']>400).sum()/ \
                    len(set_data[3].data[j]['FL1-A'])*100

    # save to csv
    percent_positive.to_csv('percent_positive_reactor'+str(i+1)+'.csv')

    plt.plot(percent_positive.index, percent_positive['percent'])
    ax.set_yscale('log')
    plt.title('Percentage of cells above threshold '+str(threshold)+
            ' reactor '+str(i+1))

    # approximate the switching rate as the slope of the linear part in the
    # log plot
    # reactor 1:
    # from 12.6 h to 17 h
    # reactor 2:
    # from 14.5 h to 18 h
    # reactor 3:
```

```
    # from 14.2 h to 18 h

    indexes=np.logical_and(percent_positive.index>=H_time_threshold[i,0],
                            percent_positive.index<=H_time_threshold[i,1])

    # add the relevant parts in bold red to the plot
    plt.plot(percent_positive.index[indexes],
                percent_positive['percent'][indexes],
                'r', linewidth=2)

    # fit a line to the data
    # use a linear regression
    slope, intercept, r_value, p_value, std_err = linregress(
        percent_positive.index[indexes],
        np.log(percent_positive['percent'][indexes]))

    print('Reactor '+str(i+1)+':')
    print('slope: '+str(slope))
    print('intercept: '+str(intercept))
    print('r_value: '+str(r_value))

    # plot the linear regression
    plt.plot(percent_positive.index[indexes],
                np.exp(intercept)*np.exp(slope*percent_positive.index[indexes]),
                'k--')

    # save the parameters
    parameters.loc[i, 'H']=slope
    parameters.loc[i, 'H_r_value']=r_value
    parameters.loc[i, 'H_r2']=r_value**2

    ## measure the maximum growth rate

    # set_data[0][0].data → OD
    fig, ax=plt.subplots()

    ax.plot(set_data[0][0].data['t'], set_data[0][0].data['m'],'r', label='OD')

    ax.legend()

    # x ax limits 0 to 25
    plt.xlim([0,25])

```

```
    # estimate growth rate between 6 and 12 hours, 12.5 h, and 12.2 h
    # by fitting an exponential to the OD data

    limits=[12,12.5,12.2]
    indexes=np.logical_and(set_data[0][0].data['t']>=6,
                           set_data[0][0].data['t']<=limits[i])

    # fit an exponential to the data
    # use a linear regression
    slope, intercept, r_value, p_value, std_err = linregress(
        set_data[0][0].data['t'][indexes],
        np.log(set_data[0][0].data['m'][indexes]))

    print("--------------------")
    print("Growth rate reactor "+str(i+1)+":")
    print('slope: '+str(slope))
    print('intercept: '+str(intercept))
    print('r_value: '+str(r_value))

    # plot the linear regression
    plt.plot(set_data[0][0].data['t'][indexes],
             np.exp(intercept)*np.exp(slope*set_data[0][0].data['t'][indexes]),
             'k--')

    # save the parameters
    parameters.loc[i, 'growth_rate']=slope
    parameters.loc[i, 'growth_rate_r_value']=r_value
    parameters.loc[i, 'growth_rate_r2']=r_value**2

parameters.to_csv('parameters.csv')

plt.show()
```

Python script for data representation:

```
from mbiomas import fca
from datetime import datetime
import matplotlib.pyplot as plt
import pandas as pd
import numpy as np

print('Reading Files in Folders')
Channel = "FITC-A"
# FCF,FCD,FCN=fca.read.multitel("batch/Reactor1",fields=['FSC-A','FSC-H',Channel],
```

```
                clean=True,zeros=True,negs=True,doublets=True,delete=True)
FCF,FCD,FCN=fca.read.multitel("continuous/Reactor1",fields=['FSC-A','FSC-H',Channel],
                clean=True,zeros=True,negs=True,doublets=True,delete=True)
Stats = fca.statistics.calculate(FCF,FCD,fields=[Channel])

#print(FCF.data["Time"])

##########################
#Plotting

#Time Scatter
fig,ax=plt.subplots(figsize = (5, 2.5))
TS=fca.plot.Plot(FCF)
TS.time_scatter(ax, FCF, FCD, y_label=Channel,y_log_lim=[1,6],  y_res=150)
plt.tick_params(axis='both', which='both', labelsize=8)
plt.plot(Stats.data[Channel]["Time"], 10**Stats.data[Channel]["Q75"],
         color = "red",linewidth=1,
        marker='o', markerfacecolor='red', markersize=3, alpha = 0.4)
plt.xlabel('Time [h]', fontsize=10)  # Adjust the fontsize as needed
plt.ylabel('FL1-A [a.u.]', fontsize=10)  # Adjust the fontsize as needed

# adjust depending on the experiment:
# plt.text(1,12000,s='D = 0.32',fontsize=14)
plt.vlines(x = 15.497,ymin=0,ymax=10**8,color="b")
# plt.text(16,12000,s='D = 0.13',fontsize=14)
plt.vlines(x = 43.73,ymin=0,ymax=10**8,color="b")
# plt.text(44,12000,s='D = 0.1',fontsize=14)
# plt.vlines(x = 91.25,ymin=0,ymax=10**8,color="b")
#plt.text(92,12000,s='D = 0.17',fontsize=14)

plt.xlim([0,90])
plt.ylim([10,20000])

fig.savefig('FL_Duplicate.png',
    bbox_inches='tight',dpi=700)
```

Additionally, the function used to show the fluxes of cells was:

```
def flux_time_scatter(ax, FCFiles,FCData, y_label='FSC-A',
                                list=None,
                                y_log_lim=None, y_res=500,
                                y_scale='log',
                                y_lin_lim=None,
                                cmap='seismic',
                                **kwargs):
```

```
    """

    Plots a scatter-plot of the fluxes of cell values across time
    for channel `index`.
    This is actually a 3D histogram, thus some of the options
        are related to bins of the histogram.

    Parameters
    ----------
    ax : matplotlib.axes.Axes
        The matplotlib Axe object on which to plot the data.
    FCFiles : mbiomas.core.mFCFiles
        The mFCFiles object containing the data.
    FCData : mbiomas.core.mFCData
        The mFCData object containing the data.
    y_label : str, optional
        The label of the y-axis.
    list : list, optional
        The list of graph from which to take space to make the colorbar,
        by default all of them.
    y_log_lim=[2,8] : array-like, optional
        A size 2 1-dimension array/array-like with the minimum and maximum
        of the y axis, in log_10. Default is based on data.
        Can only be set if `y_lin_lim` is not set.
    y_res : int, optional
        The number of bins on the y-axis. Default is 500.
    y_scale='log' : str, optional
        The scale of the y-axis. Default is 'log'. Can be 'log' for a
        log10 scale, 'symlog' for a symlog scale, or 'linear' for a linear
        scale.
    y_lin_lim : array-like, optional
        A size 2 1-dimension array/array-like with the minimum and maximum
        of the y axis, in linear space. Default is based on data.
        Can only be set if `y_log_lim` is not set.
    **kwargs : dict, optional
        Additional arguments are passed to the colorbar method.
    """

    # kwargs are passed to the colorbar

    if list is None:
        list=[ax]

    if y_log_lim is not None and y_lin_lim is not None:
        raise ValueError("Both logspace_lim and normalspace_lim are set")
```

```
    elif y_log_lim is not None:
        y_lin_lim=[10**y_log_lim[0],10**y_log_lim[1]]

    times=FCFiles.data["Time"].to_numpy()

    # need to end up with a 1D array of all the data in every
    # FCData.data[i][y_label]
    # memory allocation takes time, first, iterate to get the total
    # size of the array, then second iteration to allocate it
    # and fill it

    data_size=0
    for dataset in FCData.data.values():
        data_size+=len(dataset)

    t=np.empty(data_size,dtype=np.float16)
    d=np.empty(data_size,dtype=np.float32)
    w=np.empty(data_size,dtype=np.float32)

    i=0 # current index in full array
    for j, dataset in FCData.data.items():
        current_size=len(dataset)
        t[i:i+current_size]=times[j]
        d[i:i+current_size]=dataset[y_label]
        w[i:i+current_size]=1.0/current_size
        i+=current_size

    if y_lin_lim is None:
        y_lin_lim=[np.min(d),np.max(d)]

    # logbins = np.logspace(y_log_lim[0],y_log_lim[1],y_res)
    if y_scale=='log':

        if y_lin_lim[0]<=0:
            print("-----------")
            print("-----------")
            print("WARNING:")
            print("You were attempting to represent data down to")
            print(y_lin_lim[0])
            print("in log scale. That is impossible, use symlog scale if")
            print("you really want that. In the meantime, we set the lower")
            print("values shown to 0.1")
            print("-----------")
            print("-----------")
```

```
            y_lin_lim[0]=0.1
        y_bins=np.logspace(np.log10(y_lin_lim[0]),
                        np.log10(y_lin_lim[1]),y_res)
    elif y_scale=='linear':
        y_bins=np.linspace(y_lin_lim[0],y_lin_lim[1],y_res)
    elif y_scale=='symlog':
        y_bins=np.sinh(np.linspace(np.arcsinh(y_lin_lim[0]),
                                np.arcsinh(y_lin_lim[1]),y_res))
    else:
        raise ValueError("Value for scale not recognized")

    timebins= (times[0:-1]+times[1:])/2
    # add the two extreme bins for the time:
    timebins=np.concatenate([[2*times[0]-times[1]], timebins,
                            [1.5*times[-1]-0.5*times[-2]]])

    hist2d=np.histogram2d(t, d,
        bins=[timebins.astype('float64'), y_bins.astype('float64')],
        weights=w)
    diff= 5000*np.diff(hist2d[0], axis=0)
    # diff=5000*np.gradient(hist2d[0])[0]
    diff=gaussian_filter(diff, sigma=1.2)
    # diff[diff<0]=0

    c=cm.ScalarMappable(cmap=cmap)
    vmax=np.max(np.abs(diff))
    # hb=ax.pcolormesh(timebins.astype('float64'), y_bins.astype('float64'),
    hb=ax.pcolormesh(times, y_bins.astype('float64'),
                diff.T,
                vmin=-vmax, vmax=vmax,
                # norm='asinh',
                # norm='linear',
                norm='symlog',
                cmap=c.get_cmap())

    cb=plt.colorbar(cm.ScalarMappable(norm=hb.norm, cmap=hb.cmap),ax=list,
                    **kwargs)
```

### 6 Dataset legends

The supplementary data includes:

- The triplicate batch fermentation data used to compute most parameters.

- A duplicate experiment with multiple dilution rates to assess the instability region.
- A confirmation experiment to show the instability at 0.1 $h^{-1}$. Note that samples between 34 and 39 h are unreliable due to a technical sampling error.
